# Connection of core and tail Mediator modules restrains transcription from TFIID-dependent promoters

**DOI:** 10.1101/2021.04.05.438461

**Authors:** Moustafa M. Saleh, Célia Jeronimo, François Robert, Gabriel E. Zentner

## Abstract

The Mediator coactivator complex is divided into four modules: head, middle, tail, and kinase. Deletion of the architectural subunit Med16 separates core Mediator (cMed), comprising the head, middle, and scaffold (Med14), from the tail. However, the direct global effects of tail/cMed disconnection are unclear. We find that rapid depletion of Med16 downregulates genes that require the SAGA complex for full expression, consistent with their reported tail dependence, but also moderately overactivates TFIID-dependent genes in a manner partly dependent on the separated tail, which remains associated with upstream activating sequences. Suppression of TBP dynamics via removal of the Mot1 ATPase partially restores normal transcriptional activity to Med16-depleted cells, suggesting that cMed/tail separation results in an imbalance in the levels of PIC formation at SAGA-requiring and TFIID-dependent genes. We suggest that the preferential regulation of SAGA-requiring genes by tailed Mediator helps maintain a proper balance of transcription between these genes and those more dependent on TFIID.

## Introduction

Mediator is an evolutionarily conserved coactivator complex generally required for RNA polymerase II (RNAPII) transcription. It is composed of 25 subunits in yeast and 30 subunits in humans and is organized into four discrete structural modules: head, middle, tail and the dynamically associated kinase module [1]. The head and middle modules, in conjunction with the scaffold subunit Med14, form core Mediator (cMed), which is sufficient for Mediator function *in vitro* [2,3]. Accordingly, depletion of Med14 results in a marked global decrease of RNAPII occupancy [4] and nascent transcription [5]. Similarly, disruption of the head module via a temperature-sensitive Med17 mutation or Med17 depletion drastically reduces nascent RNA synthesis across most RNAPII-dependent genes [2,5].

In contrast to the subunits of cMed, the subunits of the tail module are not essential under normal growth conditions and mainly contribute to the regulation of inducible genes [6,7]. The main function of the tail module is as an interaction interface for numerous transcriptional activators that recruit Mediator to upstream activating sequences (UASs) in response to cellular stress or other stimuli [6,8]. Strikingly, a triad of subunits from the tail module (Med2, Med3, and Med15) can form a stable sub-complex separate from cMed when Med16 (also known as Sin4) is deleted (*med16*Δ) [9]. This stable sub-complex can bind activators *in vitro* and associates with regulatory sequences *in vivo* independent of the other modules of the Mediator complex at a handful of genes [9–11]. Like the other components of the tail module, Med16 is important for stress-induced transcription in yeast and other organisms [12–14]. However, several studies have suggested that it might have a repressive function under normal growth conditions. For instance, cells lacking Med16 show constitutive reporter expression from the *PHO5* promoter, normally induced in response to low environmental phosphate [15]; genes encoding enzymes involved in maltose metabolism [16]; *ARG1*, normally induced by isoleucine and valine starvation [9]; and *FLR1*, normally induced in response to various drugs [11]. Deletion of *MED16* also rescues expression of *HO*, encoding an endonuclease responsible for initiating mating type switching, in cells lacking various cofactors [17] or with specific mutations in the *HO* promoter [18]. On a global scale, microarray analysis has revealed substantial overlap in transcripts with increased abundance in *med16*Δ cells and cells deleted for subunits of the kinase module [6,19,20], the function of which appears to be to antagonize tail-dependent Mediator recruitment to UASs [8,21].

What is the basis of increased gene expression in the absence of Med16? One proposed model is that the separated tail enhances transcription via direct or indirect stimulation of pre-initiation complex (PIC) formation and transcription [9,11,22]. In this view, connection of the tail to cMed limits the ability of the tail triad to act as a nonspecific transcriptional activator and predicts that impairment of tail function in the absence of Med16 should suppress transcriptional overactivation. Indeed, the increased basal expression of *PHO5* observed in *med16*Δ cells can be suppressed by a mutation in Med15 [23], while deletion of *MED2* eliminates the overexpression of *FLR1* caused by *MED16* deletion [11]. Reporter expression from a mutant *HO* promoter, mediated by *med16*Δ, can likewise be negated by deletion of *MED15* [18]. Deletion of *MED2, MED3*, or *MED15* also reduces *med16*Δ-induced basal expression of *ARG1* to varying extents [9]. Lastly, mutations in *MED16* enhance transcriptional activation at a distance [24]. Relatedly, recent work indicates that the recruitment of multiple coactivators to the cell cycle-regulated *HO* promoter is enhanced in *med16*Δ strain, again suggesting that the tail/cMed connection limits the potential of the tail to facilitate transcription [18].

While there have been many studies of the role of Med16 in gene regulation, important questions remain. All of the aforementioned studies used measurement of steady-state RNA levels in *med16*Δ strains. Steady-state RNA abundances are influenced by both synthesis and decay and thus cannot be used to conclusively determine the influence of Med16 loss on transcription. This issue may be compounded by the use of deletion mutants, wherein it is difficult to disentangle the direct and indirect effects of Med16 absence. It is also unclear whether the tail triad remains globally associated with the genome when Med16 is removed. Here, we report a genome-wide analysis of the consequences of both acute and chronic Med16 loss. Through sequencing of newly synthesized RNA (nsRNA), we demonstrate that both deletion and auxin-inducible degron (AID)-mediated depletion of Med16 result in transcriptional upregulation of TFIID-dependent genes and downregulation of coactivator-redundant (CR) genes. We show that the tail triad remains globally bound to UASs in the absence of Med16, and that promoter association of cMed is decreased at downregulated genes. Disruption of the separated tail triad by co-depletion of Med15 and Med16 attenuated the transcriptional upregulation observed in Med16 depletion, arguing that the tail triad is involved in the overactivation observed with Med16 removal. Lastly, we show that co-depletion of Med16 and Mot1, a SWI/SNF-family ATPase responsible for removing TBP from TATA-containing promoters, partially rescues both the downregulation and upregulation seen with Med16 depletion alone. Taken together, our results indicate that the connection of the tail module to cMed is important for a proper balance of transcription between different classes of genes.

## Results

### Med16 removal results in moderate transcriptional overactivation

Previous studies of gene expression in *med16*Δ cells showed a dual effect: reduced expression of inducible genes upon stimulation [25,26] but increased basal expression of inducible genes [11,16,23]. However, these results are potentially confounded by indirect effects due to cellular adaptation to chronic tail separation and the measurement of steady-state mRNA levels, which are subject to transcriptional buffering effects [27]. To address the direct transcriptional effects of tail separation, we generated a yeast strain in which Med16 is tagged with an auxin-inducible degron (Med16-AID) in the presence of the auxin-interacting ubiquitin ligase component *Os*TIR1 [28]. Treatment of Med16-AID cells with the auxin indole-3-acetic acid (3-IAA) resulted in rapid depletion of Med16, with no protein detectable by western blot after 30 min (Fig. 1A). Consistent with the role of the tail in heat shock transcription [10,12,29], Med16-AID cells grown on YPD containing 3-IAA showed substantial growth impairment at 37°C (Fig. 1B).

**Figure 1.**
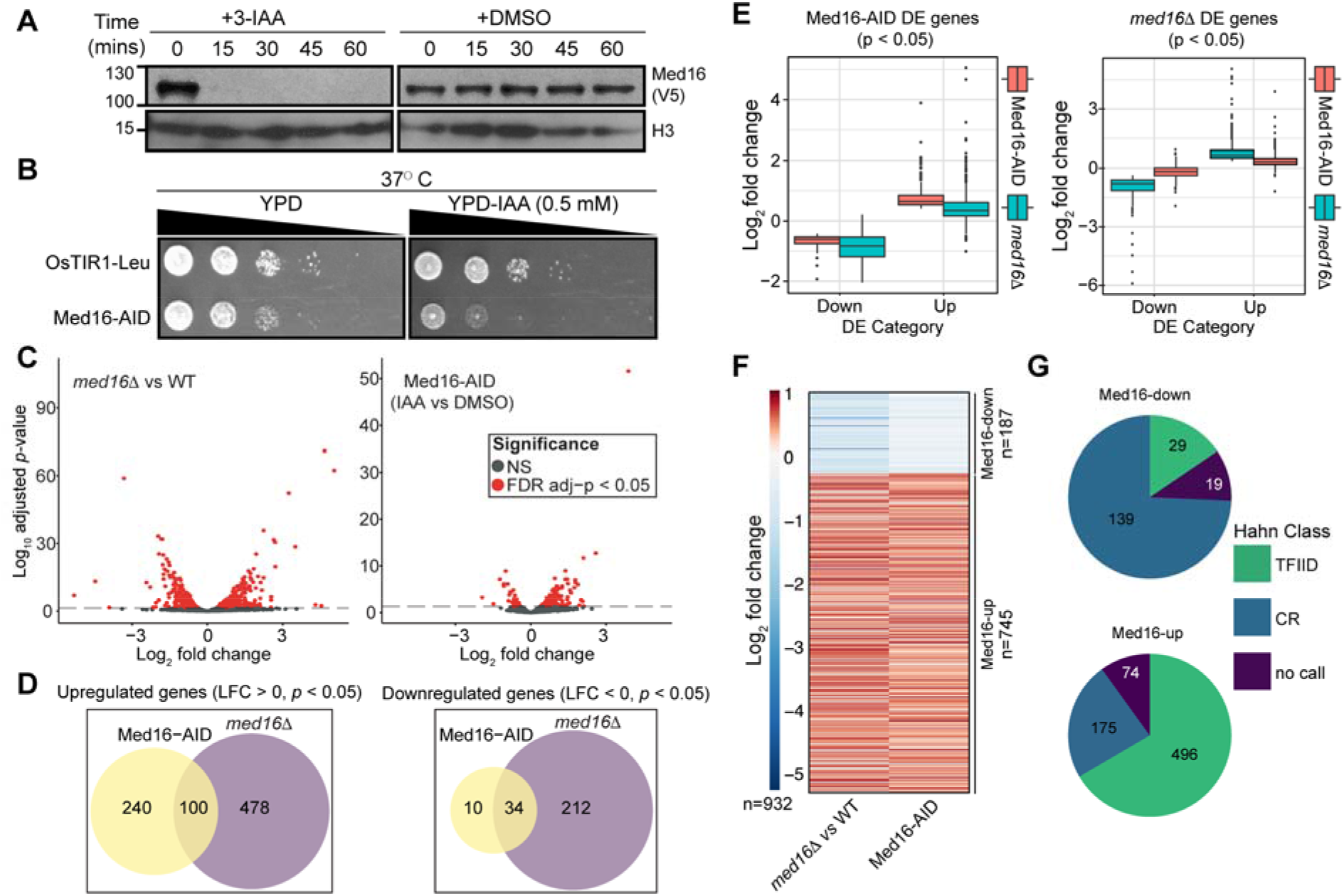
Med16 depletion primarily overactivates TFIID-dependent genes. (A) Western blot showing the kinetics of Med16-AID depletion upon 3-IAA treatment of cells. (B) Spot assays assessing growth of the parental and Med16 strains on YPD plates containing DMSO or 500 μM 3-IAA at 37°C. (C) Volcano plots of nsRNA alterations in the *med16*Δ versus WT and Med16-AID 3-IAA versus DMSO comparisons. (D) Venn diagrams of overlap in genes significantly upregulated or downregulated in the Med16-AID and *med16*Δ cells. (E) Boxplots of log_2_ fold changes in nsRNA levels for differentially expressed (DE) genes in Med16-AID (left) and *med16*Δ (right) cells. (F) *k*-means clustered (*k* = 2) heatmap of log_2_ fold changes in nsRNA levels for the set of 932 genes with concordant changes in nsRNA levels in the Med16-AID and *med16*Δ strains. (G) Pie charts indicating the classification of Med16-regulated genes according to the categories of Donczew *et al* [67].

To assess the transcriptional impact of Med16 depletion, we performed metabolic labeling of nsRNA with 4-thiouracil (4tU) followed by isolation and sequencing of nsRNA (nsRNA-seq) on Med16-AID cells after 30 minutes of 3-IAA treatment. To enable comparison of the effects of acute and chronic Med16 deficiency, we also performed nsRNA-seq in WT and *med16*Δ cells. Prior to RNA extraction, all cultures were spiked with a defined fraction of 4tU-labeled *S. pombe* cells to enable quantitative normalization. Biological replicates showed good clustering as assessed by principal component analysis (PCA) (Fig. S1A). We then performed a systematic analysis of transcriptional changes in both strains. In *med16*Δ cells, we detected 824 genes significantly changed (adjusted *p* < 0.05 by Wald test), with 578 (70.1%) upregulated (Fig. 3C). The transcriptional changes observed in Med16-AID cells were more limited, with 384 genes significantly altered and 340 (88.5%) upregulated (Fig. 1C). Of the genes upregulated in Med16-AID, 100/340 (29.4%) were shared with *med16*Δ, while 34/44 (77.3%) of Med16-AID downregulated genes overlapped with *med16*Δ (Fig. 1D). While there are far fewer genes significantly dysregulated by Med16 depletion versus deletion, we found that the genes changed in *med16*Δ were also concordantly dysregulated, albeit to a lesser extent in Med16-AID (Fig. 1E). This observation suggests that indirect effects are not a major contributor to the changes in transcript levels observed in *med16*Δ, but rather that the changes observed are direct effects whose magnitude is amplified by complete, persistent lack of Med16 versus its rapid depletion.

To focus our analysis on genes regulated similarly in both *med16*Δ and Med16-AID cells, we generated a combined list of 1,068 genes significantly altered in either strain and only retained genes whose expression was changed in the same direction in both strains. This resulted in a list of 932 genes (187 downregulated (Med16-down), 745 upregulated (Med16-up)) with consistent changes in nsRNA levels between the two strains (Fig. 1F). We then classified Med16-regulated genes according to a recent study of nascent transcription following acute depletion of SAGA and TFIID subunits, which classified genes as coactivator-redundant (CR, dependent on both SAGA and TFIID for maximal expression) or TFIID-dependent [30]. Of the 187 Med16-down genes, 168 were annotated in the prior study, and 139 (82.7%) were classified as CR, while the majority of the Med16-up genes with an annotation (496/671, 73.9%) were TFIID-dependent (Fig. 1G). The former observation is consistent with previous work indicating preferential regulation of SAGA-dominated genes by the Mediator tail [6,7,31]. Med16-down genes were also enriched for TATA boxes (Fig. S1B), consistent with the reported prevalence of TATA elements in CR promoters [30].

### The Mediator tail triad remains globally bound to UASs when severed from cMed

Previous studies have shown that, at individual loci, the tail module remains bound in *med16*Δ cells [9,11]. However, it is unclear if this is a general phenomenon or restricted to certain regions. To assess the generality of these observations, we profiled the genome-wide binding of Mediator modules in WT and *med16*Δ cells using chromatin endogenous cleavage and high-throughput sequencing (ChEC-seq), which we previously used to efficiently map Mediator binding to UASs [32–34]. We generated strains bearing MNase-tagged derivatives of the tail (Med2, Med3, Med5, Med15, and Med16 (WT only)), the head subunit Med8, the middle subunit Med9, the kinase subunit Med13, and the scaffold subunit Med14 in the WT and *med16*Δ backgrounds. We noted increases in the levels of Med2 and Med8 and a decrease in the level of Med5 in the *med16*Δ strain (Fig. S2A). We also generated a free MNase control strain in which FLAG-tagged MNase is driven by the *MED8* promoter, used in our previous studies of Mediator binding to the genome [33], was integrated at the *ura3* locus. As with Med8-MNase produced from its endogenous locus, we observed increased expression of *pMED8-*driven MNase in the *med16*Δ background (Fig. S2A). We then performed three replicate experiments for each factor. Visualization of ChEC-seq data along a representative segment of the yeast genome revealed distinct patterns of Mediator association in the *med16*Δ strain. Tail triad subunit (Med2, Med3, and Med15) occupancy was only slightly decreased, consistent with previous single-locus ChIP studies [9,11] (Fig. 2A). Med5 dissociated from the genome in the absence of Med16, consistent with biochemical analyses [35], though a caveat to this result is that we observed decreased Med5 protein levels in the *med16*Δ strain (Fig. S4B). Lastly, binding of the cMed subunits (Med8, Med9, and Med14) and the kinase module subunit Med13 was markedly impaired in *med16*Δ. We also note that, consistent with increased free MNase expression and potentially the increased chromatin accessibility of *med16*Δ cells [15,36], higher free MNase signal is present in *med16*Δ datasets.

**Figure 2.**
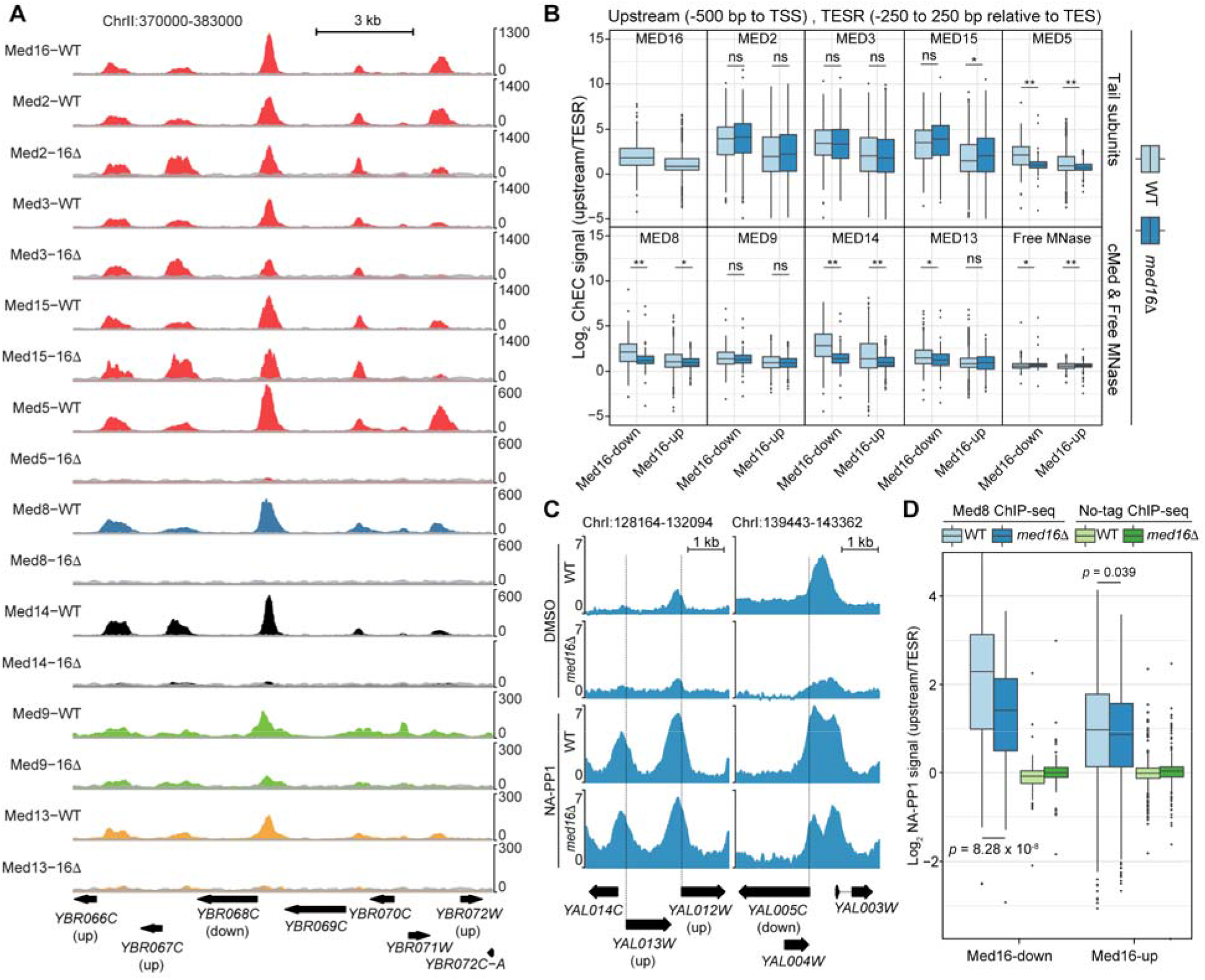
Effects of tail separation on Mediator association with the genome. (A) Tracks of Mediator tail (Med2, Med3, Med5, Med15, Med16), middle (Med9), head (Med8), kinase (Med13), and scaffold (Med14) ChEC-seq signal at a representative region of the yeast genome from WT and *med16*Δ cells. Free MNase signal from the appropriate strain is overlaid (grey) on each track for comparison. (B) Boxplots of log_2_ upstream/TESR Mediator and free MNase ChEC-seq signal from the WT and *med16*Δ strains for Med16-down and Med16-up genes. All comparisons within each cluster are by Wilcoxon rank sum test: ns indicates no significant difference, * indicates *p* < 0.01, and ** indicates *p* < 0.001. (C) Tracks of log_2_ Med8/no-tag ChIP-seq signal from the WT *kin28as* and *med16*Δ *kin28as* strains treated with DMSO or NA-PP1. (D) Boxplots of log_2_ upstream/TESR Med8 and no-tag ChIP-seq signal from the WT *kin28as* and *med16*Δ *kin28as* strains treated with NA-PP1 for Med16-down and Med16-up genes. Statistical differences between groups were assessed by Wilcoxon rank-sum test.

To more systematically assess the effects of *MED16* deletion on Mediator UAS binding, we investigated binding upstream of Med16-down and Med16-up genes. ChEC-seq replicates were highly consistent (Fig. S2B-C) and were thus averaged for this analysis. As discussed above, the levels of select Mediator subunits, as well as free MNase, are altered in *med16*Δ cells (Fig. S2A). The increased free MNase signal in *med16*Δ datasets potentially complicates between-sample comparisons, as ratios between Mediator subunit and free MNase signal may be artificially compressed due to increased negative control signal. We therefore sought a within-sample normalization strategy. We reasoned that, within a single ChEC-seq sample, comparison of Mediator-bound regions (that is, UASs) to non-Mediator-bound regions of accessible chromatin would provide a measure of the specificity of each experiment. To this end, for each gene in the Med16-down and Med16-up clusters, we determined ChEC-seq signal in a 500 bp window upstream of the TSS and divided it by the signal in a 500 bp window centered on the transcription end site region (TESR) of the same gene, as TESs are nucleosome-depleted [37–39] but would not be expected to be Mediator-occupied. As expected, we observed low enrichment of upstream signal over TES for free MNase at either gene cluster in WT and *med16*Δ cells (Fig. 2B). In concordance with single locus results, the tail triad subunits (Med2, Med3, and Med15) showed no significant reduction in upstream/TES enrichment between WT and *med16*Δ for both gene clusters. However, we noted a modest but significant increase in Med15 enrichment upstream of Med16-up genes. cMed subunits (Med8 and Med14) and Med5 showed a significant decrease in their enrichment ratios in *med16*Δ relative to WT (*p* < 0.01 by Wilcoxon rank-sum test) (Fig. 2B). On the other hand, Med9 did not show a statistically significant reduction in upstream over TES enrichment in *med16*Δ compared to WT for both gene clusters, possibly due to less efficient cutting resulting in a lower signal-to-noise ratio (Fig. 2A-B). Furthermore, the kinase subunit Med13 showed a significant reduction of upstream over TES enrichment in *med16*Δ compared to WT for cluster 1 genes but not cluster 2, which might again be reflective of reduced cutting efficiency. However, we note that it has previously been reported that nuclear depletion of Med16 has little effect on kinase module occupancy of selected loci [10].

To confirm that a 30 min 3-IAA treatment of Med16-AID cells is sufficient to yield similar effects on Mediator genomic occupancy, we performed ChEC-seq for Med2 (tail), Med14 (scaffold), and Med5 (tail) after DMSO or 3-IAA treatment. The levels of the tagged subunits did not change following Med16 depletion further supporting the specificity of the AID system (Fig. S3A). We performed the experiment in biological duplicates for each condition that showed high reproducibility (Fig. S3B-C). As observed in *med16*Δ cells, Med2 UAS occupancy was only mildly reduced by Med16 depletion. In contrast, Med5 and Med14 UAS occupancy at both Med16-regulated gene clusters was greatly reduced (Fig. S3D). Taken together, these data thus show that, on a genome-wide scale, loss of the tail/cMed connection leaves the tail triad associated with UASs while dissociating cMed subunits from these same regions.

### cMed associates with promoters in cells lacking Med16

Our ChEC-seq results indicate that *MED16* depletion results in the dissociation of cMed subunits from UASs. However, this approach does not assay Mediator binding to promoters, likely due to steric occlusion of DNA by the PIC [33], and so it is unclear how cMed/tail dissociation affects promoter association of cMed, which can occur in cells lacking multiple tail subunits [7,8,40]. Analysis of Mediator enrichment at promoters requires inhibition or depletion of the TFIIH kinase subunit Kin28, which results in impaired RNAPII CTD phosphorylation and subsequent trapping of PIC-associated Mediator [41,42]. We therefore generated WT and *med16*Δ cells bearing the *kin28as* (analog-sensitive) allele, which can be reversibly inhibited by the ATP analog NA-PP1, and an HA-tagged Mediator subunit. We tagged Med8 as a representative subunit of cMed, while Med5 was chosen because it dissociates from UASs in *med16*Δ cells and is absent from Mediator purified from *med16*Δ cells [35]. We first visually assessed Med8 binding relative to no-tag control experiments at selected regions of enrichment. Without *kin28as* inhibition, we detected reduction of Med8 signal in the *med16*Δ background, reflecting disconnection of cMed from the tail and consequent loss of UAS association (Fig. 2C). With NA-PP1 treatment, we observed a robust increase in Med8 signal, consistent with promoter trapping of cMed, with no or moderate reduction in *med16*Δ (Fig. 2C). We next analyzed Med8 and Med5 binding to the promoters Med16-down and Med16-up genes. Med8 occupancy at Med16-down promoters was significantly reduced in *med16*Δ cells, while there was minimal effect on its binding to Med16-up promoters (Fig. 2D, S4A). We were unable to detect Med5 enrichment in *med16*Δ cells (Fig. S4A). It should be noted that we detected a reduction in Med5 protein levels in the *med16*Δ strain (Fig. S4B); however, the biochemical evidence that Med16 is required for Med5 association with cMed [35] suggests that this result is not simply an artifact of altered protein levels. Taken together, these results indicate that the cMed/tail connection is involved in but is not essential for Mediator promoter recruitment, consistent with previous studies indicating dispensability of multiple tail subunits for promoter recruitment of cMed [7,8,40]. Lastly, these data provide *in vivo* support for the biochemically-defined role of Med16 in anchoring Med5 to cMed.

### Transcriptional activation following Med16 depletion is tail-dependent

To better understand the mechanism underlying the transcriptional overactivation observed upon separation of cMed and the tail module, we sought to determine its dependence on the remaining tail subunits. We depleted Med15 and Med16 simultaneously, reasoning that loss of Med15, a major point of interaction for transcription factors [43–46], would substantially impair any ability of the independent tail triad to activate transcription. We also depleted Med15 alone to enable comparison of the effects of cMed/tail separation from those of directly impairing tail function. We observed robust depletion of targeted factors in the Med15-AID and Med15/16-AID strains after 30 min of 3-IAA treatment (Fig. 3A), and so used this time point for 4tU labeling. Biological replicates showed good clustering by PCA (Fig. S5A). We first assessed the impact of Med15 and Med15/16 depletion on genes dysregulated by Med16 removal. In terms of impact on Med16-up genes, removal of Med15 had no significant impact alone or in combination with Med16 loss (*p* = 0.89 by pairwise Wilcoxon rank-sum test), though we noted a larger interquartile range (Fig. 3B). Interestingly, Med16-up genes were also upregulated by degradation of Med15 on average, though to a significantly lesser extent, and concurrent removal of Med15 and Med16 significantly reduced the upregulation observed in both the Med15-AID and Med16-AID strains (*p* < 2.2 x 10^−16^ by pairwise Wilcoxon rank-sum test for all upregulated gene comparisons) (Fig. 3B). We conclude that the transcriptional overactivation mediated by cMed/tail separation is at least partially dependent on the independent tail triad, which remains associated with the genome in the absence of Med16. Furthermore, removal of Med15 alone is sufficient to moderately increase transcription of some genes.

**Figure 3.**
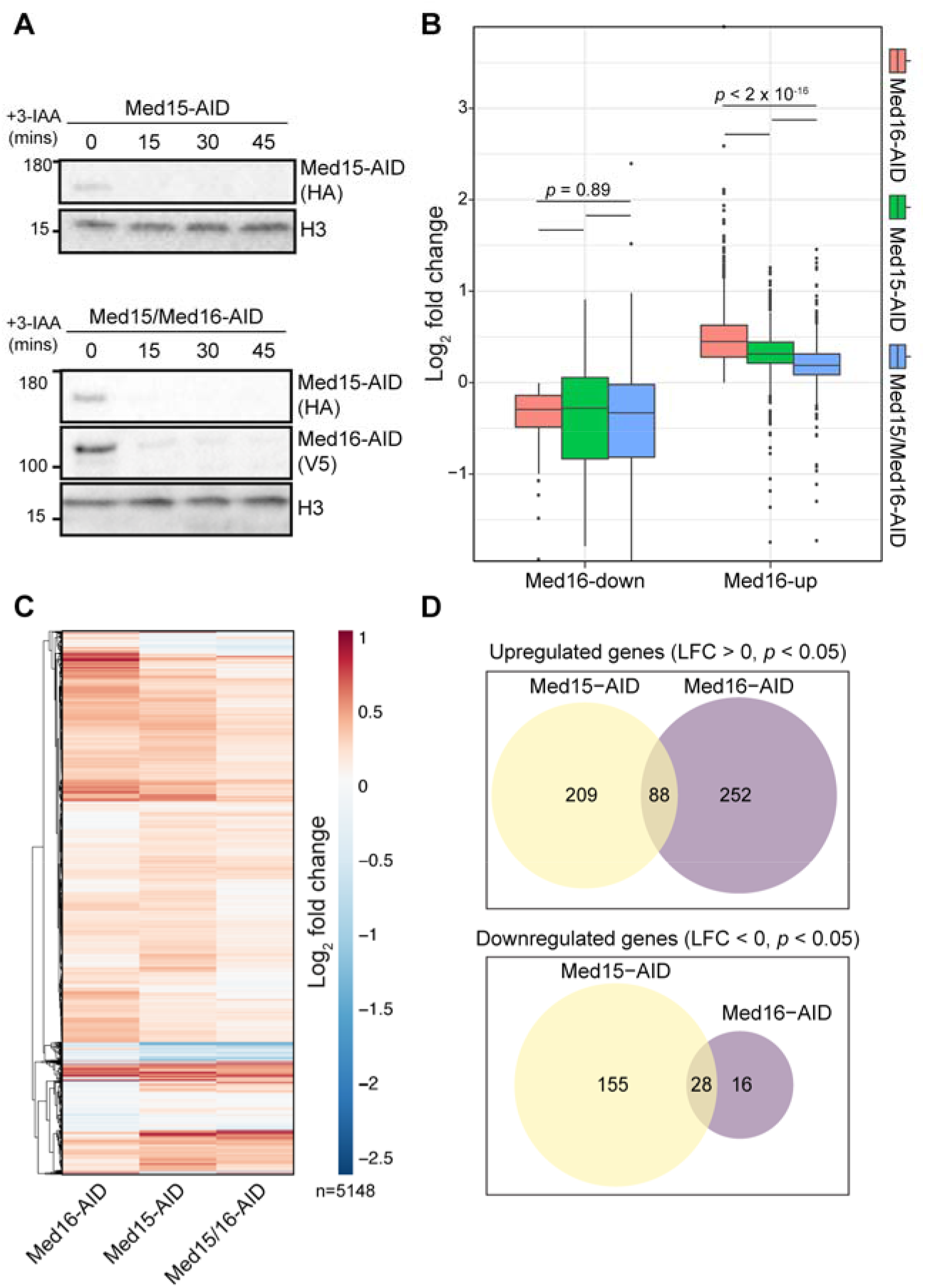
Transcription overactivation by Med16 removal is partially dependent on the tail triad. (A) Western blots showing the kinetics of Med15-AID and Med16-AID depletion upon 3-IAA treatment of single or double-degron cells. (B) Boxplots of log_2_ fold changes in nsRNA levels of transcripts produced from Med16-AID downregulated and upregulated genes for the Med15-AID, Med15/16-AID, and Med16-AID 3-IAA versus DMSO comparisons. Statistical differences between groups were assessed by Wilcoxon rank sum test with Holm correction for multiple testing. (C) Hierarchically clustered heatmap of log_2_ fold changes in nsRNA levels of transcripts produced from 5,148 genes encoding verified ORFs for the Med15-AID, Med15/16-AID, and Med16-AID 3-IAA versus DMSO comparisons. (D) Venn diagrams of overlap between genes significantly upregulated or downregulated in the Med15-AID and Med16-AID strains. Statistical differences between groups were assessed by pairwise Wilcoxon rank sum test with Holm correction for multiple testing.

We next compared global transcriptional changes in the Med15-AID, Med16-AID, and Med15/16-AID strains in order to compare the effects of cMed/tail separation and removal of a tail subunit. We plotted fold changes for 5,148 genes encoding verified ORFs as a hierarchically clustered heatmap (Fig. 3C). Here, we observed relatively similar patterns of transcriptional alterations between Med16-AID and Med15-AID, though we noted some small distinct clusters. Consistent with this, there was notable divergence in the genes significantly dysregulated by Med16 and Med15 depletion (Fig. 3D). From these analyses, we conclude that the transcriptional consequences of severing the tail from cMed are not necessarily equivalent to those of directly compromising tail function by depleting an activator-interacting subunit.

Lastly, we examined the impact of Med15, Med16, and Med15/16 depletion on the expression of CR and TFIID-dependent genes. Notably, Med16 removal did not downregulate CR genes on average (Fig. S5B), suggesting that there is instead a subset of such genes particularly sensitive to cMed/tail separation. Med15 removal resulted in a slight overall downregulation of CR genes, while co-depletion of Med15 and Med16 downregulated CR genes to a moderately greater extent than either single degron. Depletion of either Med16 or Med15 led to a modest overall upregulation of TFIID-dependent genes, which was partially suppressed by their co-depletion (Fig. S5B).

### Impaired TBP redistribution dampens the transcriptional effects of Med16 removal

A primary function of Mediator is to promote formation of the PIC [41,47,48]. Based on our data to this point, we speculate that the downregulation of Med16-down genes is due to reduced activator-dependent recruitment, reflected by lower promoter occupancy of cMed (Fig. 2C), and a consequent attenuation of PIC formation. As removal of Med15 in the context of Med16 depletion partially rescues the enhanced transcription of Med16-up genes, we surmise that the independent tail plays a role in promoting PIC formation through a means other than cMed recruitment, as tail separation has no apparent impact on the association of cMed with these promoters (Fig. 2D). To test the idea that alterations in PIC formation might underlie the transcriptional changes caused by Med16 loss, we sought a means by which to shift the balance of PIC assembly back toward a wild-type state. For this, we considered depletion of Mot1, a SWI/SNF-family ATPase that removes TBP from intrinsically favorable binding sites [49–52]. Med16-down gene promoters are enriched for TATA boxes (Fig. S1B), which are high-affinity TBP binding sites; thus, restricting TBP removal from these promoters might help restore PIC formation in the context of reduced cMed association, while making less TBP available for transcription of Med16-up genes, enriched in TFIID-dependent genes. Both Mot1 and Med16 were efficiently depleted with a 30 min 3-IAA treatment (Fig. 4A) and so we chose this time point for RNA labeling. As in previous experiments, Med16/Mot1-AID nsRNA-seq was performed in triplicate (Fig. S6A). As we hypothesized, Mot1 removal both increased the transcription of Med16-down genes (*p* = 4.84 x 10^−11^ by Wilcoxon rank-sum test) and decreased the transcription of Med16-up genes (*p* < 2.2 x 10^−16^ by Wilcoxon rank-sum test) (Fig. 4B). These observations support the idea that some degree of alteration in PIC formation underlies the transcriptional dysregulation associated with absence of Med16. On the whole-transcriptome level, we observed several gene clusters for which removal of Mot1 reversed the change in expression caused by Med16 depletion alone (Fig. S6B). We also noted clusters in which Mot1 removal attenuated or enhanced the transcriptional upregulation induced by Med16 loss.

**Figure 4.**
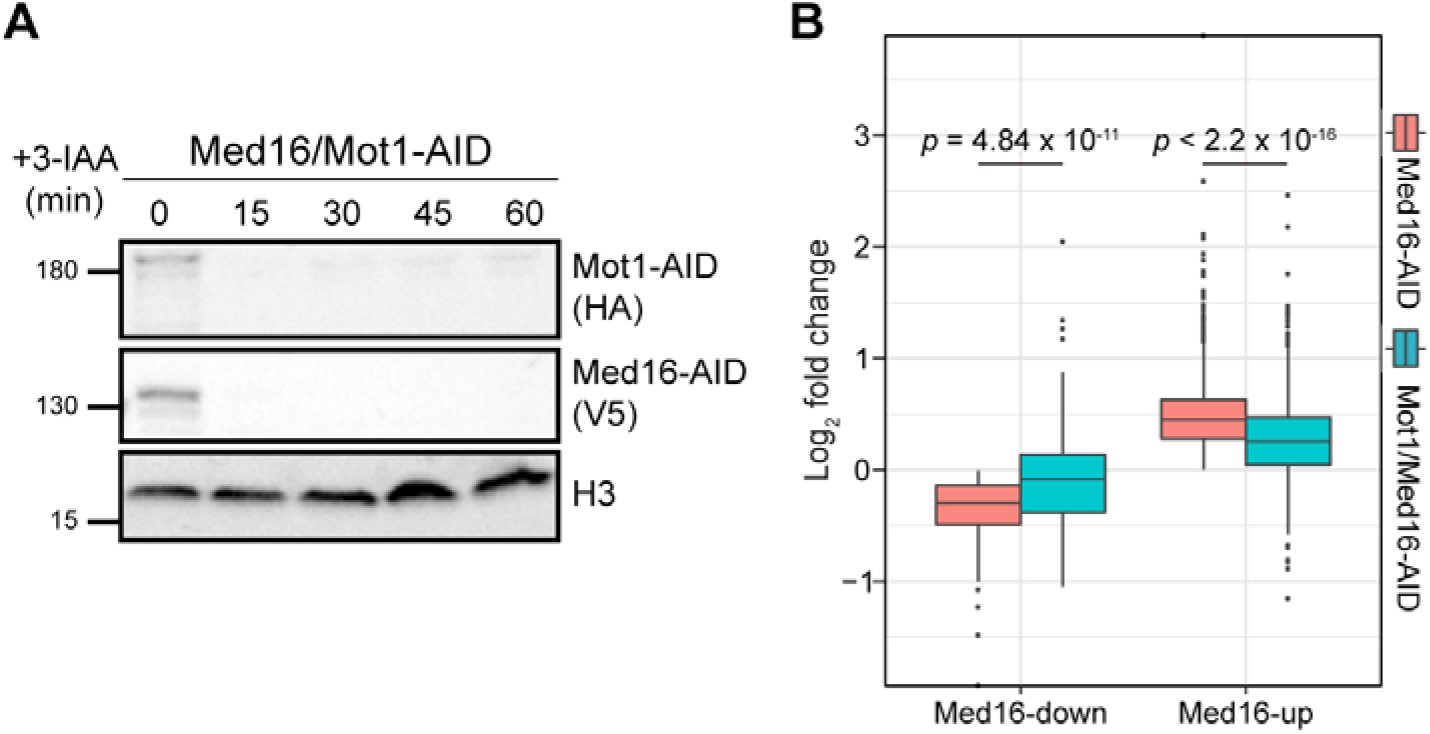
Mot1 removal partially restores wild-type transcription in Med16-AID cells. (A) Western blots showing the kinetics of Mot1-AID and Med16-AID upon 3-IAA treatment of double-degron cells. (B) Boxplots of log_2_ fold changes in nsRNA levels of transcripts produced from Med16-down and Med16-up genes for the Med16-AID and Mot1/Med16-AID 3-IAA versus DMSO comparisons. Statistical differences between groups were assessed by pairwise Wilcoxon rank sum test with Holm correction for multiple testing.

## Discussion

In this study, we confirm that Med16 depletion severs Mediator into separate cMed and tail subcomplexes and that these subcomplexes bind independently to regulatory sequences. Transcriptional downregulation in the absence of Med16 is likely to depend in part on reduced cMed recruitment and a concomitant attenuation of PIC formation, while transcriptional upregulation upon loss of Med16 appears to depend on the separated tail module and enhanced PIC formation. Increased gene expression in the absence of Med16 was previously reported [6,19,20], but the use of knockout strains and measurement of steady-state transcript levels potentially obscured the direct transcriptional effects of Med16 deficiency. We addressed these issues through a combination of acute depletion and quantification of spike-in normalized nsRNA. Our data show concordance in gene sets dysregulated by Med16 deletion and depletion, suggesting that the transcriptional effects of *MED16* deletion are mainly direct.

Activator-dependent recruitment of Mediator relies on the interaction of the tail module with activators for recruitment to UASs [7,8,40] and appears to predominate at genes regulated by SAGA (i.e., CR genes), based on the preferential impact of tail subunit deletions on the transcript levels [6,7] and TFIIB promoter occupancy [8] of SAGA-dominated genes. When cMed is disconnected from the tail by Med16 deletion or depletion, its recruitment to CR genes is compromised. Given that Mot1 removal significantly restores the transcription of genes downregulated by Med16 loss, we surmise that this reduced cMed recruitment reduces PIC formation. In this view, removal of Mot1 results in persistent TBP association with TATA boxes, which are enriched in Med16-down promoters (and CR gene promoters as a whole [30]), thus partially bypassing the requirement for full cMed recruitment in PIC formation.

At genes upregulated by cMed/tail separation, the situation is less clear. At these genes, the absence of Med16 has little effect on cMed promoter association, consistent with activator-independent recruitment via interactions with the PIC [7]. Previous studies indicated that transcriptional overactivation of selected genes in *med16*Δ cells could be suppressed by removal of Med15, a major activating-binding subunit of the tail module [18,23]. We now show that removal of Med15 strongly suppresses the global overactivation observed with depletion of Med16, indicating a role of the independent tail triad in this overactivation and generalizing the previous single-locus results. It has previously been speculated that the independent tail triad can promote PIC formation [9,11,22]. Indeed, our finding that Mot1 removal strongly suppresses Med16-depletion-dependent transcriptional overactivation suggests some involvement of increased PIC formation in this tail triad-dependent process. How might the independent tail promote PIC assembly and/or stability? Physical interactions between Med15 and TFIIE have been reported *in vivo* and *in vitro* [53–55], and so the independent tail could conceivably stabilize the PIC formed at overactivated genes, potentially in concert with cMed. This upregulation could also involve the SWI/SNF chromatin remodeling complex: *MED15* deletion abrogates the enhanced *HO* promoter binding of its catalytic subunit induced by *MED16* deletion, and deletion of Swi2/Snf2 attenuates *MED16* deletion-induced *HO* reporter expression [18].

Our observation that cMed/tail separation results in the downregulation of a relatively small number of primarily CR genes and moderate but pervasive upregulation of TFIID-dependent genes suggests that tailed Mediator plays an important role in balancing the transcriptional output of these different gene classes. In this view, activator-dependent recruitment of Mediator to CR genes via tail module interactions serves to not only promote the expression of these genes but also to restrict the expression of TFIID-dependent genes, where the activator-independent pathway of Mediator recruitment predominates. From a broader perspective, our results may indicate that gene-specific coactivator functions are important not only for appropriate expression of their direct target genes but also for restricting the expression of other large gene sets, thus helping to enforce transcriptional balance.

## Materials and Methods

### Yeast methods

Mediator subunits were tagged with 3xFLAG-MNase using pGZ109 (HIS3MX6 marker) [56] or 3xHA using pFA6a-3HA-TRP1 [57]. *MED16* was deleted from the *kin28as* strain SHY483 by replacement with the hygromycin resistance cassette from pAG32 [58]. *kin28as* strains were subsequently transformed with the Kin28-expressing plasmid pSH579 (kindly provided by Steven Hahn) to increase Kin28 protein levels as described and grown in SC-ura to maintain the plasmid. AID strains were constructed in strain SBY13674 (W303 expressing *pGPD1-OsTIR1-LEU2*, kindly provided by Sue Biggins). Med16 was tagged with 3xV5-IAA7 using pL260/pSB2065 (kanMX6 marker) [59] while Med15 and Mot1 were tagged with 3xHA-IAA7 (HIS3MX6 marker) using pGZ360. Strain genotypes are provided in Table S1.

### ChEC-seq

ChEC-seq was performed as previously described with a 1 minute calcium treatment [34]. One replicate each of Med3, Med8, and Med14 ChEC-seq in WT cells were previously published [34] (GSE112721) and reanalyzed here. ChEC-seq libraries were prepared by the Indiana University Center for Genomics and Bioinformatics (CGB) using the NEBNext Ultra II DNA Library Prep Kit for Illumina. Libraries were sequenced for 38 or 75 cycles in paired-end mode on the Illumina NextSeq 500 platform at the CGB.

### nsRNA-seq

nsRNA-seq was performed as previously described with minor modifications [30]. Briefly, cultures were treated with 3-IAA (final concentration of 0.5 mM) or DMSO for 30 min at 30°C. Following treatment, cultures were labelled for 6 min at 30°C with 4tU (final concentration of 5 mM). A spike-in of separately labeled *S. pombe* culture was then added to a final ratio of 1:4 (*S. pombe* to budding yeast). Total RNA was extracted using the Masterpure Yeast RNA extraction kit (Lucigen MPY03100) as per the manufacturer’s protocol, and subsequent biotinylation, pulldown, and purification was done as previously described [30]. rRNA was depleted from the purified nsRNA fraction using Terminator™ 5′-Phosphate-Dependent Exonuclease (Lucigen TER51020) as per the manufacturer’s protocol. rRNA-depleted nascent RNA was purified and concentrated using RNAClean XP clean beads (1.8:1 beads:sample ratio). RNA-seq libraries were prepared by the CGB using the TruSeq Stranded Total RNA kit for Illumina. Libraries were sequenced at the CGB as described above.

### X-ChIP-seq

ChIP experiments were performed in duplicate as previously described with minor modifications [8]. Briefly, ATP analog-sensitive *kin28as* strains were pre-cultured in yeast nitrogen base (YNB) medium lacking uracil before inoculation in yeast extract-peptone-dextrose (YPD) medium. Cultures were treated with 6μM of 1-Naphthyl PP1 (NA-PP1; Tocris Bioscience) or DMSO for 15 min at 30°C prior to crosslinking. For each ChIP, 3 µg of mouse monoclonal anti-HA antibody (Santa Cruz Biotechnology, sc-7392) were coupled to Dynabeads coated with Pan Mouse IgG antibodies (Thermo Fisher Scientific, 11042). ChIP-seq libraries were prepared as described [60] and sequenced for 50 cycles in paired-end mode on the Illumina HiSeq 4000 platform at the McGill University and Génome Québec Innovation Centre.

## Data analysis

### nsRNA-seq

Paired-end reads were mapped to the sacCer3 (budding yeast) and ASM294 (fission yeast) genomes using STAR [65]. Read counts per gene were determined using the “--quantMode GeneCounts’’ option of STAR. Spike-in normalization and differential expression analysis were performed in R with DESeq2 [66]. For spike-in normalization, *S. pombe* read counts were used to determine library size factors using the estimateSizeFactors function. Each size factor was then used as the size factors for the corresponding *S. cerevisiae* sample prior to differential expression analysis. Results from all DESeq2 comparisons are provided in Table S2 and an annotated list of Med16-regulated genes is provide in Table S3. Heatmaps were generated using the R pheatmap package.

### ChEC-seq

paired-end reads were mapped to the sacCer3 genome build using Bowtie2 with the default settings in addition to “--no-unal --dovetail –no-discordant --no-mixed.” SAM files generated by Bowtie2 were used to create tag directories with HOMER [61]. BedGraph files were generated with the HOMER makeUCSCfile command, normalizing to a total of 1 million reads. Bigwig files were made using the “bedGraphToBigWig” program [62]. Bigwigs were visualized with the Gviz Bioconductor package [63]. Heatmaps were made using deepTools with a bin size of 10 bp in a 2 kb region centered around the TSSs of genes dysregulated by Med16 deletion or depletion. The inputs for the heatmaps were bigwig files of pooled biological replicates. For correlation analysis, total normalized ChEC-seq signal in 2 kb windows centered on the TSSs of all genes in sacCer3 (6672 genes) was determined using HOMER annotatepeaks.pl. Spearman correlation plots were created using the corrplot R package. For boxplots of all ChEC-seq experiments performed in the WT and *med16*Δ strains, we determined normalized counts in the upstream region (defined as 500 bp upstream of the annotated TSS) and the TES region (TESR, defined as the 500 bp centered on the annotated TES) and divided upstream by TESR counts to determine fold enrichment. This normalization was performed due to increased free MNase expression and potentially increased chromatin accessibility of the *med16*Δ strain relative to WT [15,36].

### ChIP-seq

Reads were aligned as described for ChEC-seq. BAM files were made using SAMtools. Bigwig files were generated from BAM files with deepTools using either SES normalization with the corresponding no-tag control sample as input or CPM normalization, both at a bin size of 10 bp [64]. Gene tracks were prepared as described for ChEC-seq using SES-normalized bigwigs as input. For boxplots, upstream/TESR normalization was performed as described above.

## Supporting information

Figures S1-S6, Legends for Tables S1-S3

## Data availability

All datasets generated in this work have been deposited in GEO (ChEC-seq and nsRNA-seq: GSE169748; ChIP-seq: GSEqqqqqqq).

## Acknowledgements

We thank David Stillman and Bobby Yarrington for valuable discussions and yeast strains.

