## Supplementary material for "Connection of core and tail Mediator modules restrains transcription from TFIID-dependent promoters": Figures S1-S6, Legends for Tables S1-S3

Contents:

Figures S1-S6
Legends for Tables S1-S3

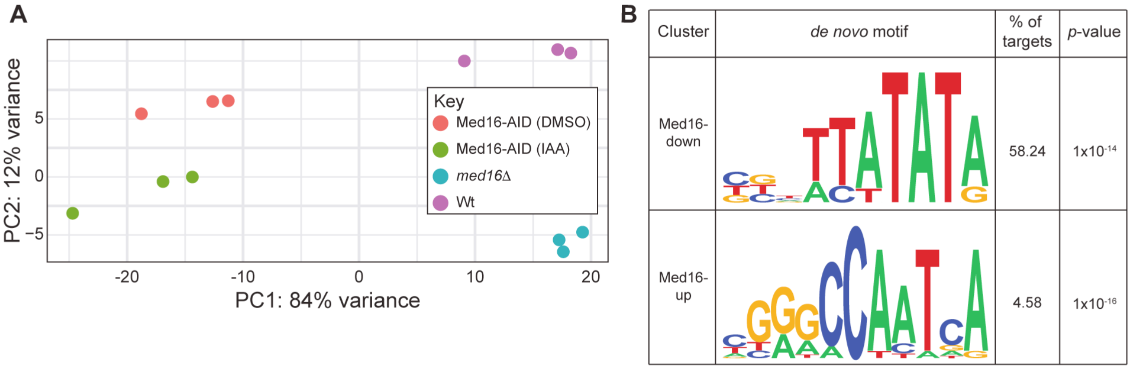


**Figure S1.**
PCA plot of replicate nsRNA-seq experiments performed in WT and *med16Δ* cells and Med16-AID cells treated with DMSO or 3-IAA. (B) Sequence logos of the *de novo* motifs discovered in the promoters (-400 to +100 bp relative to TSS) of genes in Med16R clusters.

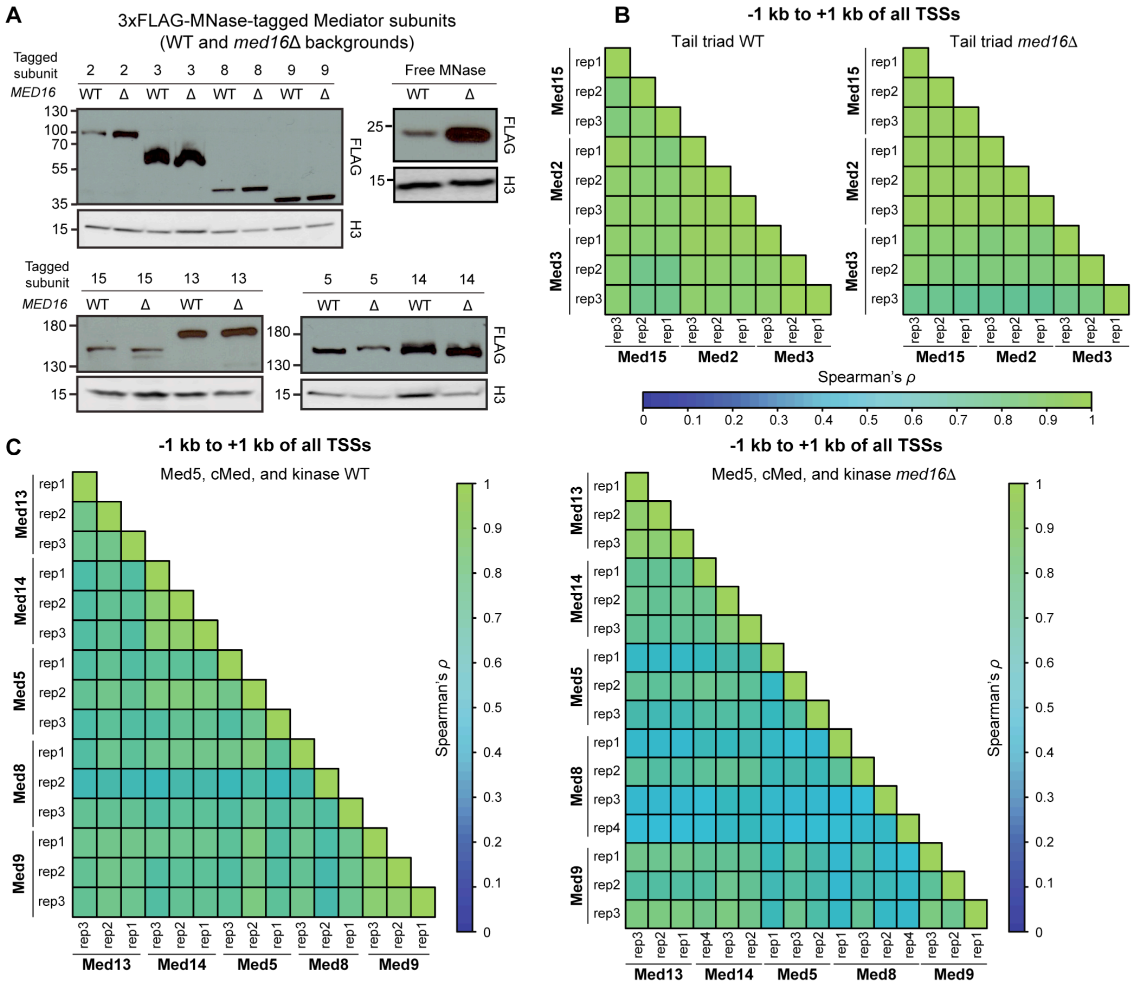


**Figure S2.**

(A) Western blots for 3xFLAG-MNase-tagged Mediator subunits in the WT and *med16Δ* strains. (B) Correlation matrices of tail triad subunit ChEC-seq replicate signal from the WT and *med16Δ* strains (-1 kb to +1 kb relative to the TSSs of all genes). (C) Correlation matrices of cMed, kinase, and Med5 ChEC-seq replicates from the WT and *med16Δ* strains (-1Kb to +1Kb relative to TSS of all genes).

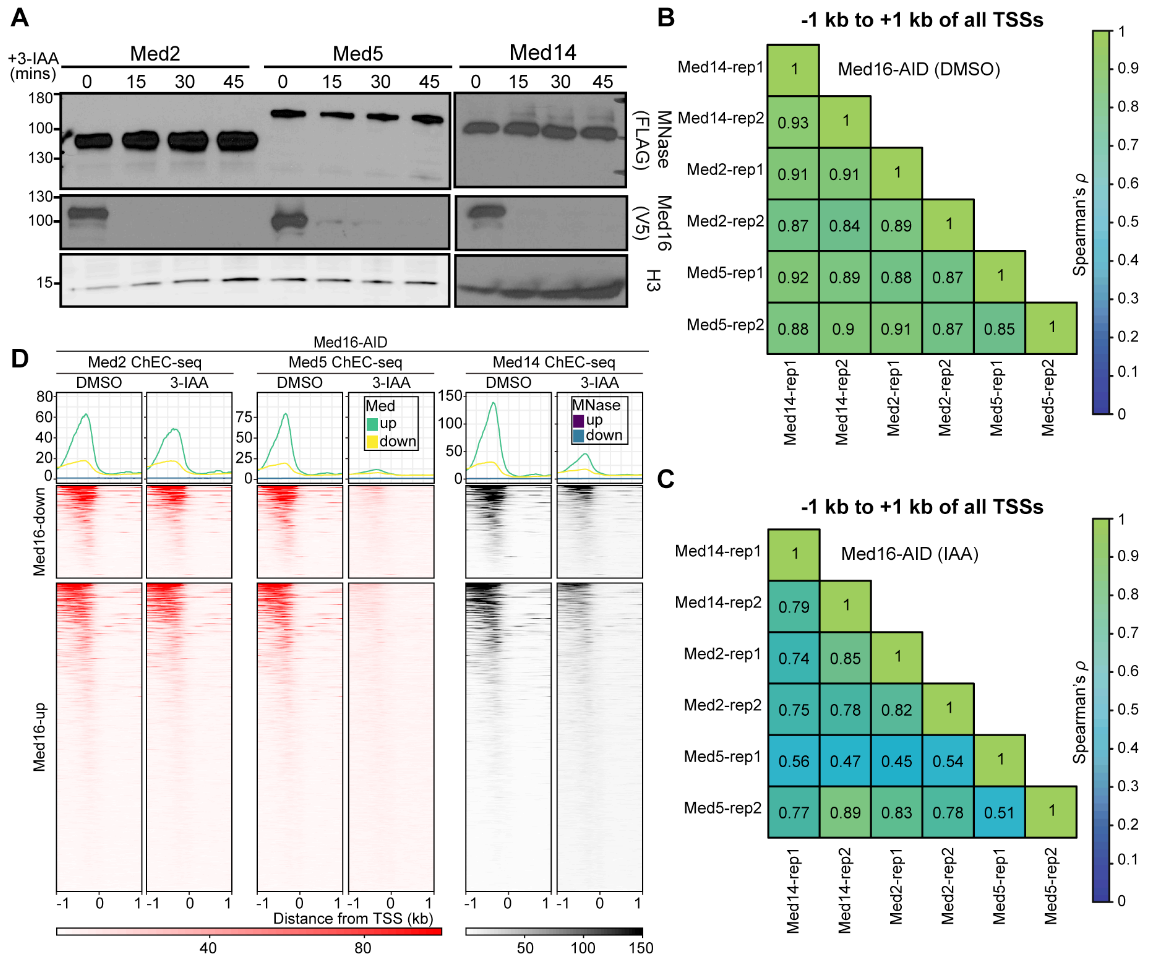


**Figure S3.**
(A) Western blots for 3xFLAG-MNase tagged Mediator subunits after treatment of Med16-AID cells with 3-IAA. (B-C) Correlation matrices of ChEC-seq replicates from DMSO- and 3-IAA-treated Med16-AID cells (-1 kb to +1 kb relative to the TSSs of all genes). (D) Heatmaps of Mediator ChEC-seq signal from Med16-AID cells treated with DMSO or 3-IAA for downregulated and upregulated genes (-1 kb to +1 kb relative to TSSs).

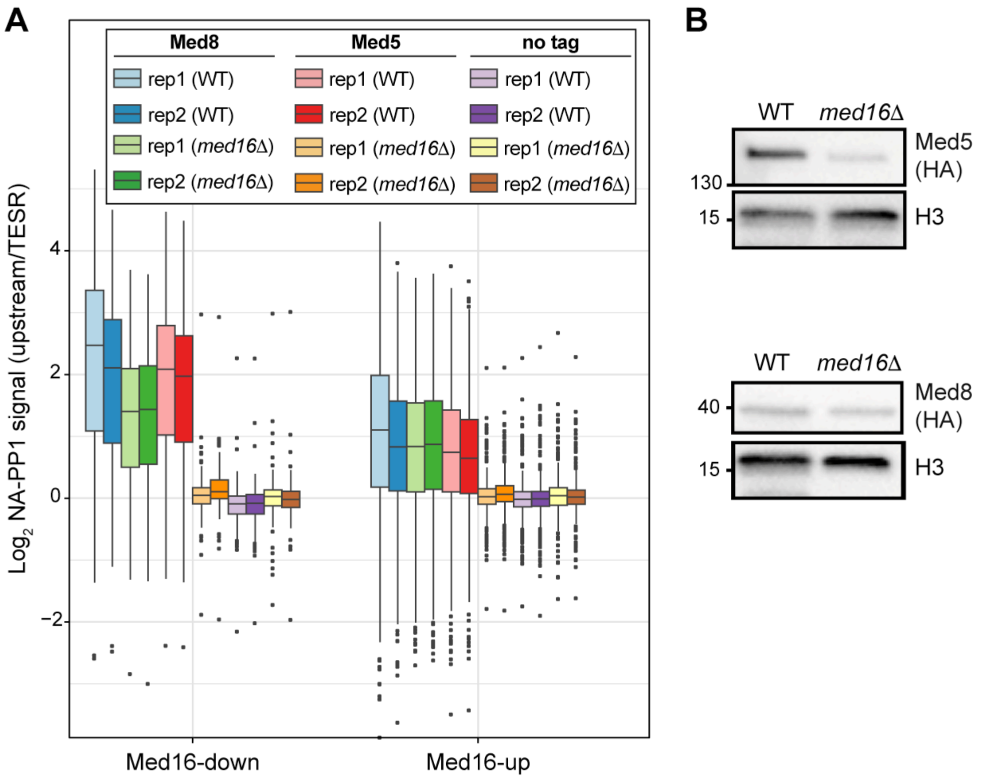


**Figure S4.**
(A) Boxplots of replicate $\log_2$ upstream/TESR Med8, Med5, and no-tag ChIP-seq signal from the WT *kin28as* and *med16Δ kin28as* strains treated with NA-PP1 for downregulated and upregulated genes. (B) Western blots for 3xHA tagged Mediator subunits in the WT and *med16Δ* strains.

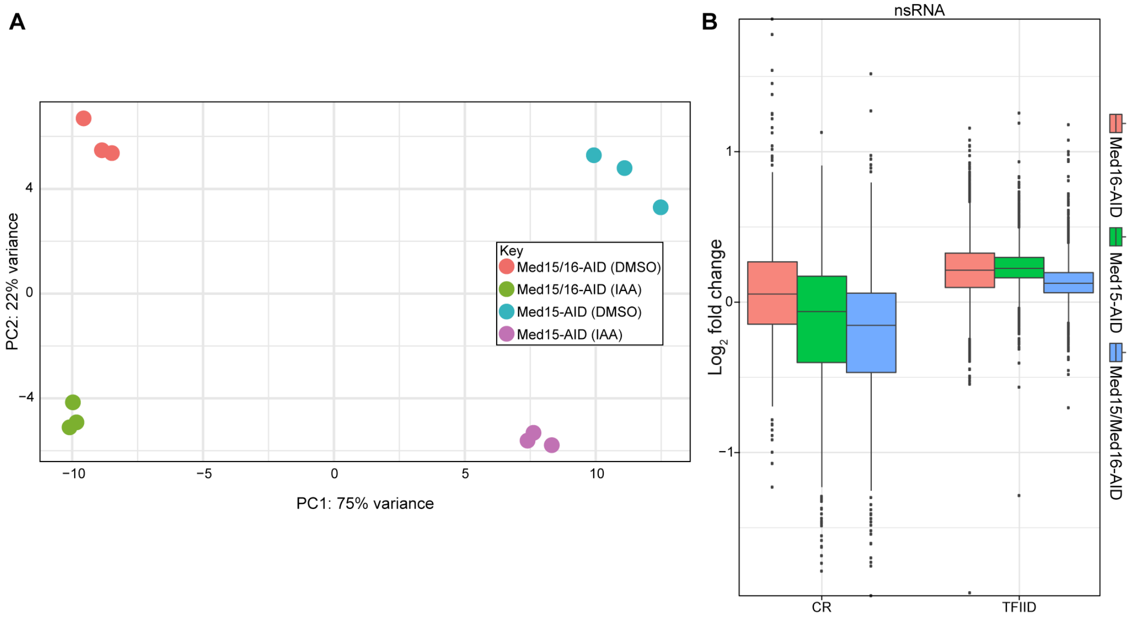


**Figure S5.**
(A) PCA plot of replicate nsRNA-seq experiments performed in Med15-AID and Med15/16-AID cells treated with DMSO or 3-IAA. (B) Boxplots of $\log_2$ fold changes in nsRNA levels of transcripts produced from CR and TFIID genes for the Med15-AID, Med15/16-AID, and Med16-AID 3-IAA versus DMSO comparisons.

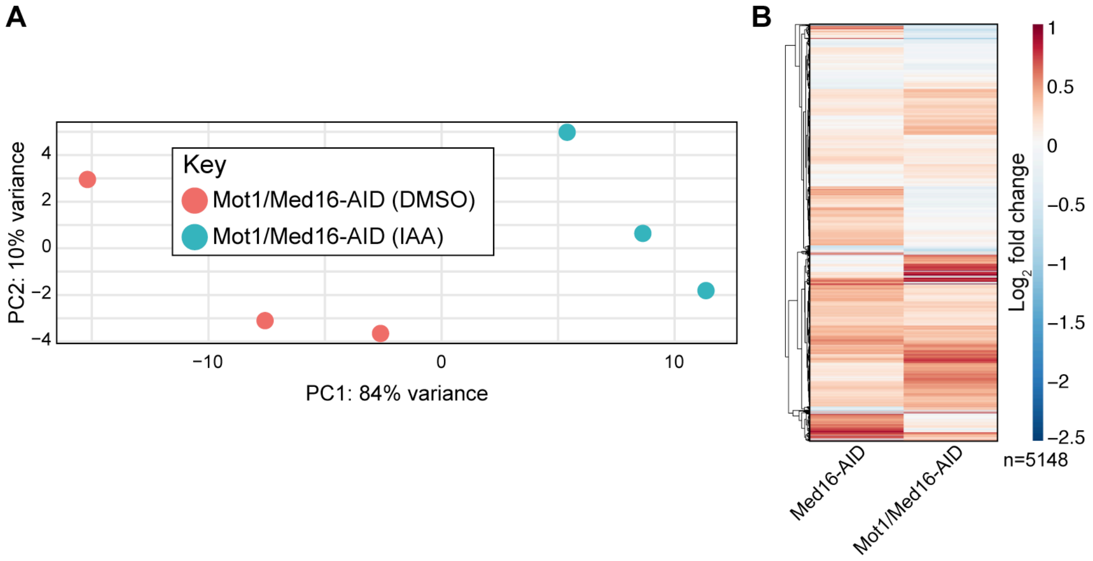


**Figure S6.**
(A) PCA plot of replicate nsRNA-seq experiments performed in Mot1/Med16-AID cells treated with DMSO or 3-IAA. (B) Hierarchically clustered heatmap of $\log_2$ fold changes in nsRNA levels of transcripts produced from 5,148 genes encoding verified ORFs for the Med16-AID and Mot1/Med16-AID 3-IAA versus DMSO comparisons.

**Supplementary Table Legends**

**Table S1.** Yeast strains used in this work.

**Table S2.** DESeq2 output for all nsRNA-seq experiments reported in this work.

**Table S3.** Annotation information for Med16-regulated genes.
